# GERV: A Statistical Method for Generative Evaluation of Regulatory Variants for Transcription Factor Binding

**DOI:** 10.1101/017392

**Authors:** Haoyang Zeng, Tatsunori Hashimoto, Daniel D Kang, David K Gifford

## Abstract

The majority of disease-associated variants identified in genome-wide association studies (GWAS) reside in noncoding regions of the genome with regulatory roles. Thus being able to interpret the functional consequence of a variant is essential for identifying causal variants in the analysis of GWAS studies. We present GERV (Generative Evaluation of Regulatory Variants), a novel computational method for predicting regulatory variants that affect transcription factor binding. GERV learns a k-mer based generative model of transcription factor binding from ChIP-seq and DNase-seq data, and scores variants by computing the change of predicted ChIP-seq reads between the reference and alternate allele. The k-mers learned by GERV capture more sequence determinants of transcription factor binding than a motif-based approach alone, including both a transcription factor’s canonical motif as well as associated co-factor motifs. We show that GERV outperforms existing methods in predicting SNPs associated with allele-specific binding. GERV correctly predicts a validated causal variant among linked SNPs, and prioritizes the variants previously reported to modulate the binding of FOXA1 in breast cancer cell lines. Thus, GERV provides a powerful approach for functionally annotating and prioritizing causal variants for experimental follow-up analysis. The implementation of GERV and related data are available at http://gerv.csail.mit.edu/

## 1 Introduction

Genome-Wide Association Studies (GWAS) have revealed genetic polymorphisms that are strongly associated with complex traits and diseases ([21, 20, 27, 12]). Missense and nonsense variants that occur in protein coding sequences are simple to characterize. However, many GWAS detected variants reside in non-coding regions with regulatory function ([12, 7]). The influence of non-coding variation on gene expression and other cellular functions is not well understood. Previous work has observed that non-coding DNA changes in the recognition sequences of transcription factors can affect gene expression and cellular phenotypes ([31]). Thus predicting the effect of genomic variants on TF binding is an essential part of interpreting the role of non-coding variants in pathogenesis. Most of existing computational approaches to predict the effect of SNPs on TF binding such as sTRAP and HaplogReg are based on quantifying the difference between the presented reference and alternate alleles in the context of canonical TF binding motifs ([1, 17, 19, 24, 30, 22, 28]). Recent work ([14]) uses k-mer weights learned from a gapped-kmer SVM ([9]) to score the effect of variants, taking into account the frequency of k-mer occurrences but not the spatial effect of k-mers.

Here we present GERV (Generative Evaluation of Regulatory Variants), a novel computational model that learns the spatial effect of k-mers on TF binding *de novo* from whole-genome ChIP-seq and DNase-seq data, and scores variants by the change in predicted ChIP-seq read counts between the reference and alternate alleles. GERV improves on existing models in three ways. First, GERV doesn’t assume the existence of a canonical TF binding motif. Instead it models transcription factor binding by learning the effects of specific k-mers on observed binding. This allows GERV to capture more subtle sequence features underlying transcription factor binding including non-canonical motifs. Second, GERV accounts for the spatial effect of k-mers and learns the effect of cis-regulatory regions outside of the canonical TF motif. This enables us to model the role of important auxiliary sequences in transcription factor binding, such as cofactors. Third, GERV incorporates chromatin openness information as a covariate in the model which boosts the accuracy of the predicted functional consequence of a variant.

We first demonstrate the power of GERV on the ChIP-seq data for transcription factor NF-*κ*B. We show that GERV learns a vocabulary of k-mers that accurately predicts held-out NF-*κ*B ChIP-seq data and captures the canonical NF-*κ*B motifs as well as associated sequences such as known co-factors. Applying GERV to six transcription factors on which allele-specific binding (ASB) analysis is available, we show GERV outperforms existing approaches in prioritizing SNPs associated with allele-specific binding. We demonstrate the application of GERV in post-GWAS analysis by scoring risk-associated SNPs and their linked SNPs for breast cancer, and show that GERV trained on FOXA1 ChIP-seq data achieves superior performance in prioritizing SNPs previously reported to modulate FOXA1 binding in breast cancer cell lines.

## 2 Methods

### 2.1 GERV Model Overview

GERV is a fully generative model of ChIP-seq reads. We assume that the genome is a long regulatory sequence containing k-mer “code words” that induce invariant spatial effects on proximal transcription factor binding. We use the level of chromatin openness in a region as a functional prior to predict the magnitude of a sequence-induced binding signal. Following this assumption, we model the read counts produced by transcription factor ChIP-seq at a given base as the log-linear combination of the DNase-seq signal on nearby bases and the spatial effect of a set of learned k-mers whose effect range covers that base.

The GERV procedure of variant scoring consists of the following three steps: (Figure 1)

1. GERV first learns the spatial effect of all k-mers (k=1 to 8) and the covariate coefficient of local DNase-seq signal over a spatial window of ± 200 base pairs (bp) *de novo* from ChIP-seq data using regularized Poisson regression
2. GERV then computes the predicted ChIP-seq read counts for the reference and alternate allele of a variant from the log-linear combination of the local DNase-seq signal and spatial effect of the learned k-mers.
3. GERV predicts the effect of a genomic variant on transcription factor binding by the *l*^2^-norm of the change of predicted reads between two alleles

**Figure 1:**
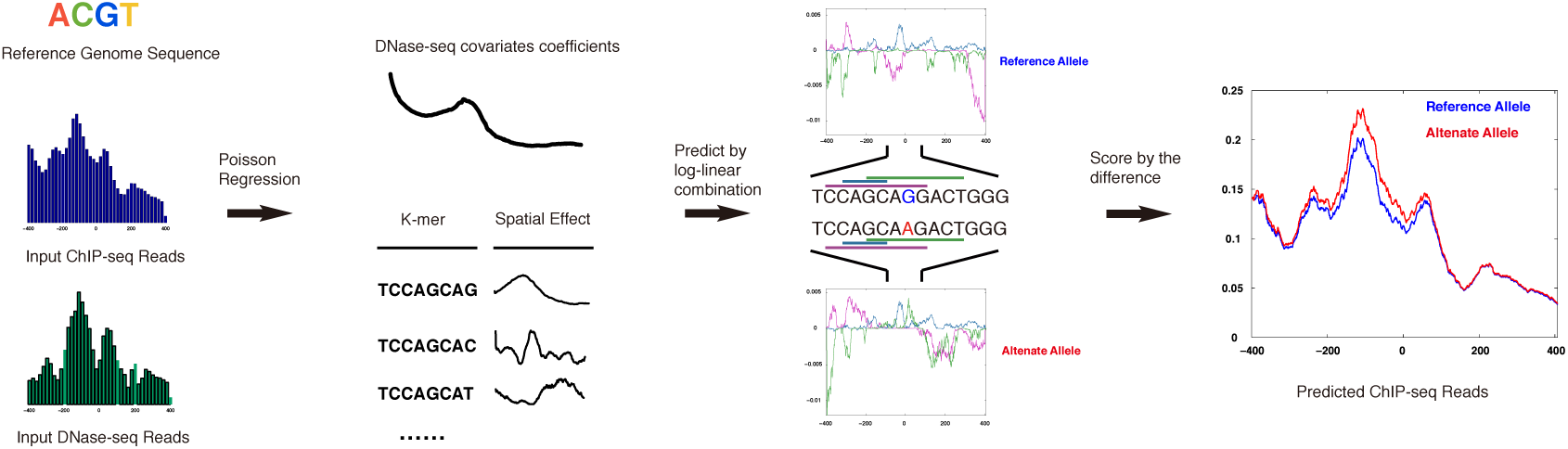
The schematic of GERV. The spatial effects of all the k-mers and the coefficients for DNase-seq covariates are learned from the reference genome sequence and ChIP-seq, DNase-seq datasets. Then the spatial effects (purple, cyan and green) of the k-mers underlying the reference (blue) and alternate (red) allele for a variant are aggregated with DNase-seq covariates by log-linear combination to yield a spatial prediction of local ChIP-seq reads for the two alleles. GERV scores the variant by the *l*^2^-norm of the predicted change of reads.

### 2.2 Learning the Spatial Effect of K-mers

The effect profile of a k-mer is defined as a real-valued vector of length 2*M* that corresponds to a spatial window of [− *M, M* − 1] relative to the start position of the k-mer. Specifically, the *j*-th entry of the profile for a k-mer is the expected log-change in read counts at the *j*-th base relative to the start of the k-mer. Here we consider k-mers with *k* from 1 to 8 (*k*_*max*_ = 8) as this is the maximum that would fit in memory in an Amazon EC2 c3.8 xlarge instance. Larger K-mers tested on a larger memory machine did not perform substantially better than 8-mers. As ChIP-seq signals are relatively sparse and spikey, we chose an effect range of ±200 bp for each k-mer (*M* = 200).

For notational convenience we use *i* for genomic coordinate, *k* for k-mer length, and *j* for coordinate offset from the start of a k-mer. We assume that the genome consists of one large chromosome with coordinate 0 to *N*. In practice we will construct this by concatenating chromosomes with the telomeres acting as a spacer. We represent the effect vector of all k-mers of length *k* as a parameter matrix *θ*^*k*^ of size 4^*k*^ × 2*M*. For any particular k-mer of length *k* starting at base *i* on the reference genome, we define 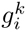 as its row index in *θ*^*k*^. So 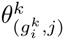 would denote the effect of this kmer at offset *j* ∈ [*−M, M −*1]. Additionally, a special parameter *θ*_0_ is used to set the average read rate of the genome globally.

The DNase-seq covariate *κ* is defined as a vector of length *N* that is aligned with each base of the genome, and we assume that ChIP-seq reads can be predicted with this covariate and the contributions from surrounding k-mers. The regression coefficients for the covariate are defined as *β* and have length 2*L*. The regression coefficients *β* can be thought of as analogous to the k-mer effect *θ*, but occurring everywhere, and scaled by the covariate *κ*. In all the experiments in this analysis, we chose an *L* = 200 to balance between computational complexity and prediction power.

Given these definitions, we define a generative model for ChIP-seq reads on the genome. Observed counts at position *i* on the genome are generated from a Poisson distribution with rate parameter *λ*_*i*_ which is defined as:

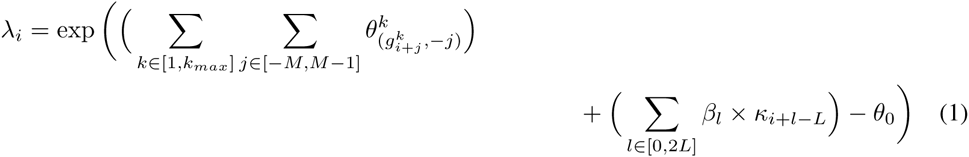

The problem we solve is a regularized Poisson regression. Particularly, we would like to maximize the following:

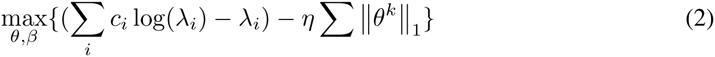

To efficiently optimize this objective function, we performed an accelerated gradient descend method. The detail of implementation can be found in the supplementary data (Supplementary Text S1).

### 2.3 ChIP-seq Signal Prediction for Reference and Alternate Allele

In step 2, given the effect profile of all the k-mers and coefficients of the DNase-seq covariate trained from step 1, we first predict the ChIP-seq count *λ* at each position across the reference genome by combining the effect of proximal k-mers and DNase-seq level into the log-linear model using equation 1. Then in similar manner, we predict the read counts *λ*^*′*^ of the alternate allele after replacing the k-mers that are affected by the variant. If we assume a Single Nucleotide Polymorphism (SNP), at most 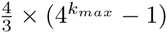 k-mers will change.

### 2.4 Variant Scoring

In step 3, we score a SNP at locus on the genome by the square root of the sum of squared per-base change (*l* ^2^-norm of the change) of binding signal at all bases within the effect range of any k-mers affected by the variant:

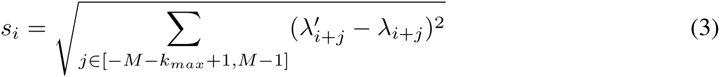

### 2.5 Collapsing GERV k-mers into a position weight matrix

We interpret the active k-mers captured by GERV with a post-processing framework that aggregates similar k-mers into position weight matrixes (PWMs):

1. We filter k-mers based on the sum of spatial effect to eliminate inactive k-mers.
2. We calculate the Levenshtein distance (number of single character edits) between the remaining k-mers.
3. We perform UPGMA hierarchical clustering over the candidate k-mers until the minimal distance among clusters is larger than 2.
4. For each cluster, we define its key k-mer as the one with the largest sum of spatial effect. We obtain the position weight matrix for this cluster by aligning all k-mers in the cluster against the key k-mer.
5. All the clusters are ranked by the average sum of spatial effect of all the k-mers in the cluster.

### 2.6 ChIP-seq Peak Prediction Comparison

Gapped-kmer SVM was downloaded from http://www.beerlab.org/gkmsvm/index.html. To match with the training data for GERV, the positive training set for gapped-kmer SVM consists of the all the NF-*κ*B ChIP-seq peaks on chr1-13 of GM12878 from ENCODE, and the negative training set consists of the same number of randomly sampled regions of similar size on chr1-13. The default parameter set (“-d 3”) was used. Both GERV and gapped-kmer SVM were evaluated on the same test set. The positive test set consists of all the NF-*κ*B ChIP-seq peaks on chr14-22 of GM12878 from ENCODE, and the negative test set consists of the same number of randomly sampled regions of similar size on chr14-22.

### 2.7 Benchmark the performance in prioritizing SNPs with allele-specific binding

#### 2.7.1 deltaSVM

deltaSVM source code was downloaded from http://www.beerlab.org/deltasvm/. For each transcription factor included in the benchmarking, a gapped-kmer SVM model was trained using ChIP-seq peaks of that factor on chr1-13 of GM12878 from ENCODE as positive sets and the same number of randomly sampled region of similar size on chr1-13 as negative sets. The default parameter set (“-d 3”) was used. As instructed by the software, the gapped-kmer SVM model was then used to score all the possible 10-mers, the result of which was input as the kmer-weight parameter to deltaSVM.

#### 2.7.2 sTRAP

We used the R version of sTRAP downloaded from the website (http://trap.molgen.mpg.de/download/TRAP_R_package) for scalability. The built-in JASPAR and TRANSFAC motif data included in the package were used. Specifically, MA0105.1, MA0105.2, MA0105.3, MA0107.1, MA0061.1, V$NFKAPPAB 01, V$NFKB Q6, V$NFKAPPAB65 01, V$NFKAPPAB50 01, V$P50 Q6, V$NFKB C and V$RELA Q6 were used for NF-*κ*B. MA0139.1, MA0531.1, V$CTCF 01, V$CTCF 02 were used for CTCF. MA0099.1, MA0099.2, MA0476.1 and V$CFOS Q6 were used for FOS. MA0059.1, MA0058.1, MA0058.2, PB0043.1, PB00147.1, V$MAX 01, V$MAX 04, V$MAX Q6, V$MYCMAX 01, V$MYCMAX 02, V$MYCMAX 03 and V$MYCMAX_B were used for MAX. MA0059.1, MA0147.1,MA0147.2, V$CMYC 01, V$CMYC 02, V$MYC 01, V$MYCMAX 01, V$MYCMAX 02, V$MYC MAX 03 and V$MYCMAX B were used for MYC. None of the JUND motifs were included in the built-in motif database of sTRAP. For each variant, the scores from different matrices of the same factor were combined by taking the highest one.

## 3 Materials

### 3.1 ChIP-seq Data

ChIP-seq data for all the factors used in this analysis were downloaded from ENCODE. The full list of GEO accession numbers can be found in Supplementary Table S1.

### 3.2 DNase-seq Data

DNase-seq data of GM12878 were downloaded from ENCODE (GEO accession GSM816665)

### 3.3 Allele-Specific Binding (ASB) SNPs

As a gold standard for SNPs that affect TF binding, we used the list of SNPs that are reported to induce allele-specific binding (ASB) of NF-*κ*B, CTCF, FOS, JUND, MAX and MYC in GM12878. The NF-*κ*B ASB SNPs are collected from [25] and [13]. The ASB SNPs data for all other transcription factors are collected from [25]. The ASB SNPs were further filtered on allele frequency (≥ 0.01) to keep only the common SNPs.

## 4 Results

### 4.1 GERV learns a vocabulary of k-mers that regulate factor binding

We first tested if GERV could predict held-out ChIP-seq data. We trained a GERV model on ENCODE NF-*κ*B ChIP-seq data and DNase-seq data from chromosomes 1-13 of GM12878, and compared the predicted ChIP-seq signal from GERV to actual ChIP-seq reads on the held-out chromosomes 14-22. The predicted ChIP-seq signals are very similar to actual ChIP-seq reads (Figure 2A,B), with a chromosome-wide Pearson’s correlation of 0.76. We measured correlation after smoothing predicted and actual reads over 400 bp windows since actual reads are insufficiently sampled to produce base-pair resolution measurements. To further examine the ability of GERV to model ChIP-seq peaks, we used the GERV model trained above to score a positive set of regions defined as all the ENCODE GM12878 NF-*κ*B ChIP-seq peaks on chr14-22, and a negative set of regions defined as same number of randomly sampled region of similar length on chr14-22. Each region was scored by the sum of predicted signal in the region. We compared GERV with a previously published kmer-based model for TF peak prediction by training a gapped-kmer SVM ([9]) on ENCODE NF-*κ*B peaks and same number of randomly sampled region of similar length on chr1-13 of GM12878, and then performing the same scoring task on the same positive and negative set. We quantified the performance of these two models in prioritizing positive regions over negative regions by calculating the area under receiver operating characteristic curve (AUROC)(Figure 2C). Our model achieved a better AUROC of 0.972 than that of 0.949 for gapped-kmer SVM. Thus GERV learns a vocabulary of k-mers that can accurately predict the ChIP-seq data.

**Figure 2:**
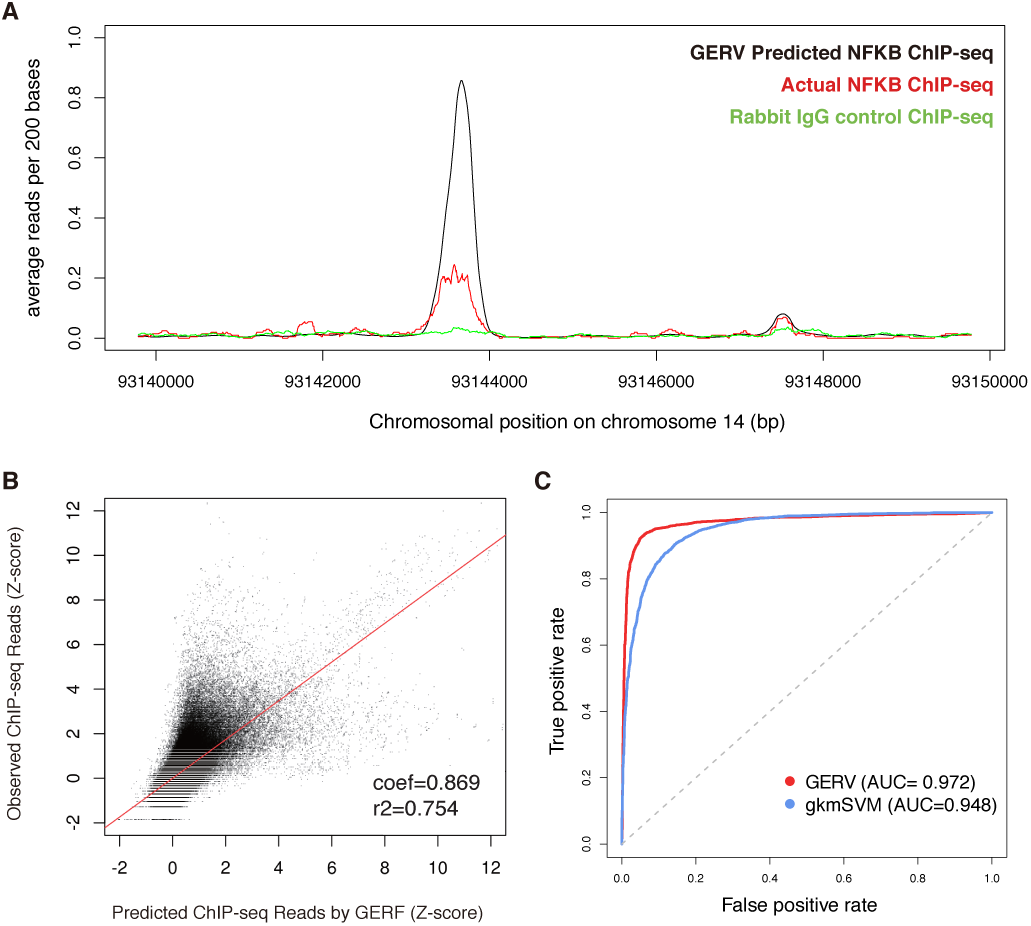
(A) Example held-out genomic region on chromosome 14 showing GERV-predicted NF-*κ*B reads (black), actual NF-*κ*B ChIP-seq reads (red), and rabbit IgG control ChIP-seq reads (green). (B) Comparison of GERV-predicted (x-axis) and observed (y-axis) NF-*κ*B ChIP-seq reads in binned regions of held-out chromosome 14-22. The coefficient and *r* ^2^ of a linear regression on predicted and actual z-score is plotted. (C) ROC curve for discriminating NF-*κ*B peaks from negative control sets using GERV and gapped-kmer SVM (gkmSVM).

Although GERV fits a model with a potentially large parameter space (± 200 bp window for 87380 k-mers when *k*_*max*_ = 8), it uses sparsifying regularization to avoid overfitting and to limit the number of active k-mers (Equation 2). For example, in the NF-*κ*B GERV model, most of the *l* ^1^-norm of the parameter matrix is contained in the top 1% of the 87380 k-mers (Supplementary Figure S1). GERV is also robust to the choice of the window size for a k-mer’s spatial effect and DNase-seq covaraites (Supplementary Table S2).

### 4.2 GERV captures the binding sequence of a TF and its co-factors

We then examined if GERV learned the sequence features important for transcription factor binding. We trained a GERV model on DNase-seq data and NF-*κ*B ChIP-seq data combined from 10 LCL individuals. Position weight matrices were generated for visualization purposes by hierarchical clustering of the active k-mers in GERV (Section 2.5) and matched to known TF motifs in JASPAR and TRANSFAC using STAMP ([18]). With a threshold of significant matching at 1e-7, many clusters of the active k-mers correspond to known motifs (Table 1). The top two k-mer clusters for NF-*κ*B were matched to motifs from NF-*κ*B family (Supplementary Figure S2A), indicating that GERV correctly learned the strongest expected sequence features for the binding. Moreover, many of the other k-mer clusters learned by GERV correspond to transcription factors which have been associated with NF-*κ*B regulation (Supplementary Figure S2B), including ETS1, AP1, IRF1 and SP1 ([26, 8, 2, 29]). To validate the role of these transcription factors in NF-*κ*B binding, we performed co-factor analysis on the same NF-*κ*B data using GEM ([10]) to search for transcription factors that have spatially binding constraint with NF-*κ*B. This analysis identified AP-1 and IRF1 as the strongest co-factors of NF-*κ*B binding. Interestingly, some of the active-kmer clusters in GERV were matched to transcription factors such as ELF1, ERF2, CTCF and SUT1 which have not been associated with NF-*κ*B binding in previous studies.

**Table 1:** TF motifs matched to active-kmer clusters in NF-*κ*B GERV model using STAMP with E-value cutoff of 1e-07. For each cluster, only the strongest match in each motif database (TRANSFAC and JASPAR) is shown. PWMs are ordered by the average sum of spatial effect of all the k-mers in the corresponding cluster.

| Cluster | Matched Motif | Motif Database | Matched TF | E-value |
| --- | --- | --- | --- | --- |
| PWM1 | M00053<br>MA0101.1 | TRANSFAC<br>JASPAR | REL<br>REL | 5.1842e-08<br>1.2145e-09 |
| PWM2 | M00053<br>MA0101.1 | TRANSFAC<br>JASPAR | REL<br>REL | 7.0388e-14<br>1.1385e-12 |
| PWM3 | M00495<br>MA0099.2 | TRANSFAC<br>JASPAR | Bach1<br>AP1 | 8.0813e-13<br>9.4186e-10 |
| PWM4 | M01111 | TRANSFAC | RBP-Jkappa | 6.0791e-08 |
| PWM5 | M00339<br>MA0080.2 | TRANSFAC<br>JASPAR | ETS1<br>SPI1 | 1.3508e-11<br>3.6387e-10 |
| PWM7 | M01057<br>MA0123.1 | TRANSFAC<br>JASPAR | ERF2<br>abi4 | 2.0724e-08<br>1.9650e-08 |
| PWM12 | MA0139.1 | JASPAR | CTCF | 1.1289e-08 |
| PWM15 | M00062<br>MA0050.1 | TRANSFAC<br>JASPAR | IRF1<br>IRF1 | 2.7655e-10<br>3.1444e-09 |
| PWM18 | MA0399.1 | JASPAR | SUT1 | 2.7097e-08 |
| PWM20 | M00722 | TRANSFAC | core-binding | 3.6831e-09 |
| PWM22 | MA0242.1 | JASPAR | run_Bgb | 3.1286e-11 |
| PWM23 | M01066 | TRANSFAC | BLIMP1 | 1.4886e-09 |
| PWM27 | MA0453.1 | JASPAR | nub | 4.2762e-09 |
| PWM32 | M00345 | TRANSFAC | GAMYB | 8.8633e-08 |
| PWM33 | MA0344.1 | JASPAR | NHP10 | 2.3002e-09 |
| PWM38 | MA0403.1 | JASPAR | TBF1 | 5.6499e-08 |
| PWM43 | M00181 | TRANSFAC | E2 | 2.9261e-08 |
| PWM49 | MA0152.1 | JASPAR | NFATC2 | 5.3905e-11 |

To further interpret the role of the transcription factors whose motifs were matched to an active-kmer clusters in the NF-*κ*B GERV model, we performed motif analysis on the SNPs known to alter transcription factor binding. Allele-specific binding (ASB) studies have identified SNPs associated with significantly imbalanced binding events on heterozygous sites ([25, 13]). Therefore we col-lected a list of 56 ASB SNPs for NF-*κ*B, and use HaploReg ([31]) to query for the motifs that these ASB SNPs altered (Supplementary Table S3). Among the 56 ASB SNPs tested, only 16 (29%) were found to alter the canonical motif of NF-*κ*B, while another 11 (20%) were found to alter the TF motif matched to other active-kmer clusters in the GERV model. Thus, GERV captures the sequence context of factor binding, which provides additional descriptive power and biological insight for auxiliary elements in TF binding.

### 4.3 GERV outperforms existing approaches in prioritizing ASB SNPs

To demonstrate the power of GERV in detecting regulatory variants, we compared GERV’s performance against existing approaches in discriminating ASB SNPs from negative control variants. We collected ASB SNPs with known differential binding for NF-*κ*B, CTCF, JUND, MAX, MYC and FOS from previous studies ([25, 13]) as positive sets, resulting in a total of 56 SNPs for NF-*κ*B, 1225 SNPs for CTCF, 26 SNPs for FOX, 233 SNPs for JUND, 71 SNPs for MAX and 69 SNPs for MYC (Section 3.3). Note that these ASB SNPs were completely held-out in the training process of any model compared in this analysis, and were only used as the test set.

For each of the six transcription factors, we constructed two types of negative SNP sets that we assume do not exhibit differential factor binding. Both kinds of negative sets are subsets of 1000 Genome Project (1KG) common (minor allele frequency *≥*1%) SNPs. In the first case we randomly sampled 100 negative samples for each positive sample, to get a reasonable sample of the background while making analyses computationally tractable. The second set is a fine-mapping task which is an important topic in post-GWAS analysis where a list of lead SNPs and their linked SNPs are under interrogation for regulatory consequence. To simulate such tasks, this second set was constructed as random selection of 1KG common SNPs within 10kb from any ASB SNP. To reflect the number of SNPs typically in a single LD block, we calculated LD information from phased genotype data in the 1KG pilot release using PLINK ([23]). With a *r* ^2^ cutoff of 0.8, the median number of linked SNPs for a variant is 10 (Supplementary Figure S3). Thus in this set we sampled 10 negative samples for each positive sample. For both types of negative sets, we sampled 10 sets with replacement so that we could obtain the mean and confidence intervals. For each of the 10 negative sets, we constructed a paired positive set, same size as the corresponding ASB SNP set, by sampling with replacement from the ASB SNPs.

For each transcription factor, we evaluated the performance of GERV as well as two published regulatory variant scoring methods sTRAP ([19]) (motif-based), and deltaSVM ([14]) (kmer-based) in discriminating the positive set from each of the two negative sets. The other motif-based methods are not included due to either the inability to produce numerical scores for the queried variants, or the low throughput that can’t scale up to thousands of SNPs. For each factor, a GERV model was trained on ENCODE ChIP-seq data from chr1-13 of GM12878, and a deltaSVM model was trained on ENCODE ChIP-seq peaks and same number of random regions of similar length on chr1-13 of GM12878. The built-in JASPAR and TRANSFAC motif dataset was used for sTRAP, which includes the motif for all the factors but JUND (Section 2.7).

We show the averaged receiver operating characteristic (ROC) curves and precision recall curves (PRC) (Supplementary Figure S4 for the first control set, Figure 3 for the second control set) of all the methods for different transcription factors and negative sets. We evaluated two aspects of the curves. The first metric is the area under curve (AUC) (Supplementary Table S4) which summarizes the overall performance in prioritizing the positive set over negative set. The second metric is the true positive rate at low false positive rate (for ROC) or the recall at high precision (for PRC), which reflects the practical need for low false discovery rate in post-GWAS analysis where thousands of lead and linked SNPs are tested for regulatory consequence. The ROC curves for GERV consistently dominated the competing methods for all factors and control scenarios, with much better AUC and higher true positive rate at low false positive rates. In PR curves because of the small size of the positive set, the confidence intervals of precision when the recall is low tend to be large, making the left-most part of the curves less informative for comparison. For transcription factor FOS, MAX and MYC, GERV achieved a PRC curve clearly superior to the others, without overlapping in the confidence interval. For factor JUND and NF-*κ*B, GERV had a similarly precision for low recall, but outperformed the other methods with consistently high precision for larger recall. For CTCF, the competing methods achieved higher precision when recalling less than 10% of the positives, but their precision dropped dramatically afterwards resulting in much lower AUC than that of GERV. Given the fact that CTCF has a motif (19 bp) more than twice as long as the maximum length of k-mer (8 bp) learnable for GERV (Section 2.2), the competitive performance on CTCF demonstrates the strong descriptive power of GERV in modeling TF binding.

**Figure 3:**
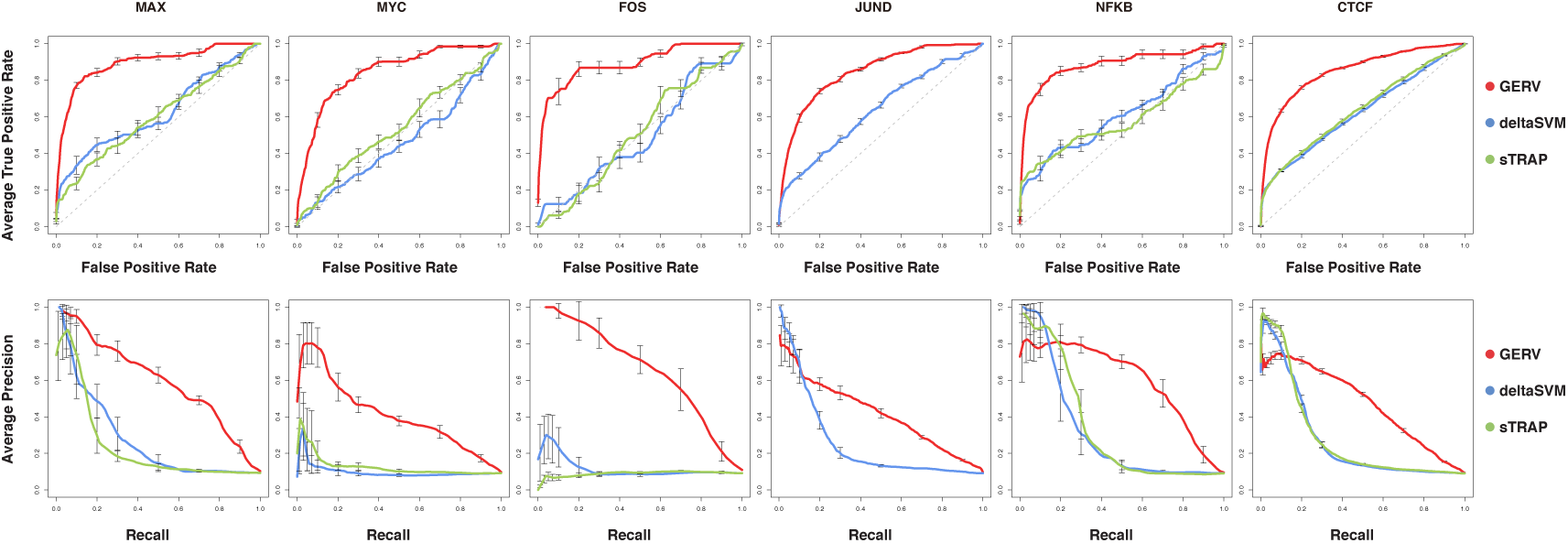
ROC (first row) and PRC (second row) curves for discriminating ASB SNPs from the second type of negative variant set (10 times of the size of positive set) using GERV (red), deltaSVM (blue), and sTRAP (green). Grey dashed line in ROC curves indicates random chance. In each figure, 95% confidence intervals of the true positive rate (for ROC) or precision (for PRC) are plotted. The performance of sTRAP on JUND is not measurable as JUND motif is not included in its built-in motif database.

We found that for our second negative control setup, choosing 50 instead of 10 negative SNPs for each positive SNP caused a noticeable decline in the area under precision recall curves (Supplementary Figure S5). With this ratio of positive to negative SNPs, deltaSVM outperformed GERV at the 10% recall point for JUND, NF-*κ*B, and CTCF, but by the 30% recall point GERV produced better precision-recall for these and all other factors.

### 4.4 GERV prioritizes linked-SNPs that modulate FOXA1 binding in breast cancer

To demonstrate the application of GERV in post-GWAS analysis, we applied GERV to a breast cancer associated variant set (AVS) collected from a previous study ([5]). It is composed of 44 risk-associated SNPs and 1,053 linked SNPs that are in strong LD with any risk-associated SNP. It has been shown that breast cancer associated SNPs are enriched for the binding sites of FOXA1, a pioneer transcription factor essential for chromatin opening and nucleosome positioning favorable to transcription factor recruitment ([11, 6, 3, 4, 16]). We trained a GERV model and a deltaSVM model on ENCODE FOXA1 ChIP-seq data from a breast cancer cell line T47D. The rs4784227 breast cancer associated SNP has been shown to disrupt the binding of FOXA1 with several lines of evidence ([5, 15]). GERV correctly predicted the effect of rs4784227 on FOXA1 binding among its linked SNPs, while deltaSVM failed (Figure 4A). Having probed a single risk-associated SNP, we then applied GERV to all the SNPs in the breast cancer AVS. The 28 variants previously reported to modulate FOXA1 binding ([5]) had significantly higher GERV scores than the rest of the AVS (Figure 4B, Mann-Whitney U test P=0.001, AUC=0.69). In contrast, deltaSVM couldn’t distinguish the positive set from the rest of the AVS (Mann-Whitney U test P=0.22, AUC=0.57)

**Figure 4:**
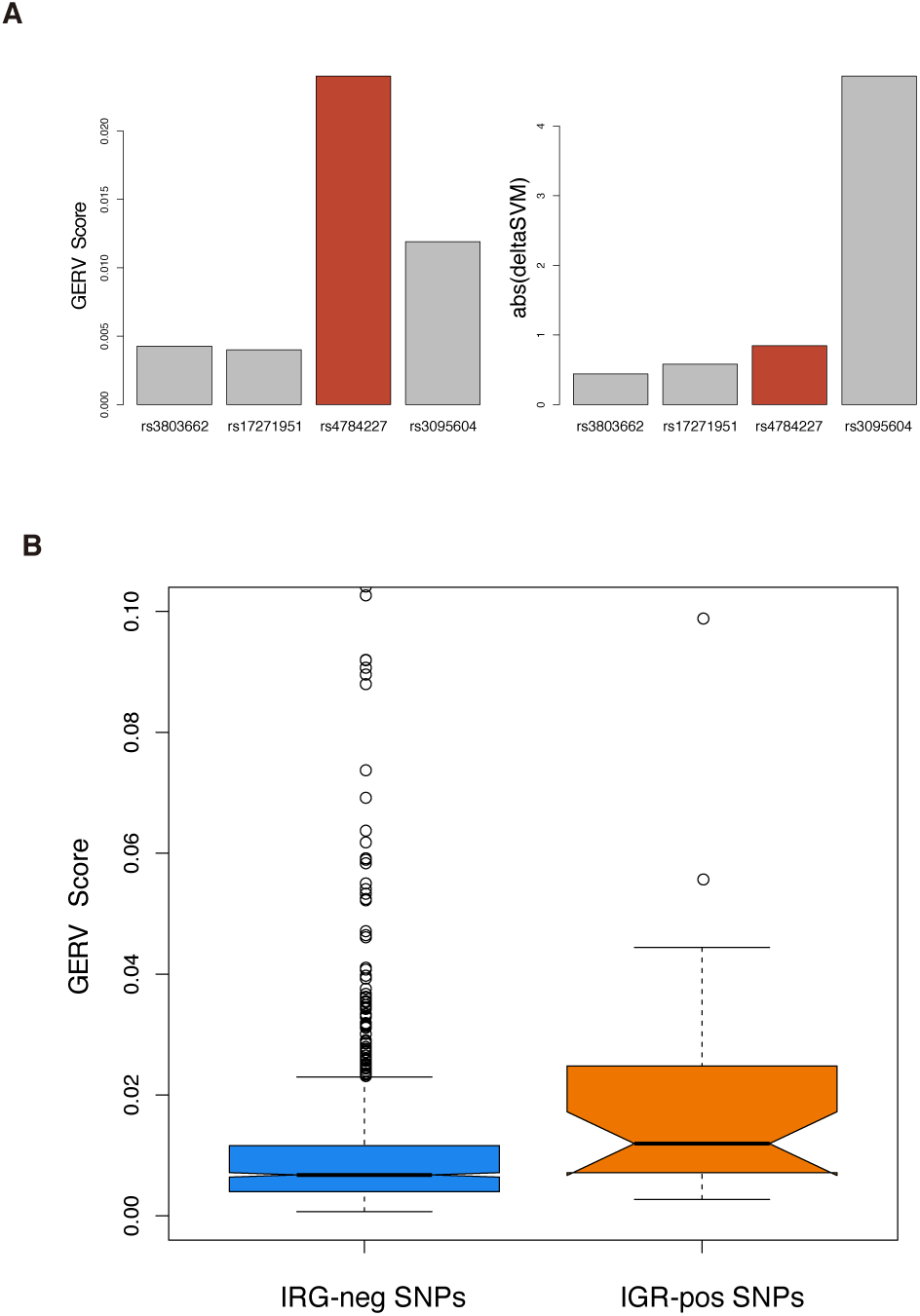
(A) GERV correctly predicted the effect of validated causal SNP rs4784227 on FOXA1 binding, while deltaSVM failed. (B) The 28 variants previously reported to modulate FOXA1 binding had significantly higher (Mann-Whitney U test P=0.001) GERV scores than the rest of the AVS (n=1065 after filtering for common SNPs)

## 5 Discussion

Despite the recent substantial advances in characterizing the genome-wide transcription factor binding sites with ChIP-seq experiments, it remains a challenge to interpret variation in the noncoding region of the genome and to determine variants that cause transcription factor binding changes in post-GWAS analysis. Our work improves the prediction of causal non-coding variants when compared to other contemporary methods.

As the first generative model that directly predicts the ChIP-seq signal, GERV achieved greater accuracy than other methods in predicting ChIP-seq peaks. GERV models the spatial effect of all the k-mers and thus captures the effect of the primary motif and auxiliary sequences on TF binding. We have shown that many of these auxiliary sequences correspond to known binding cofactors, while others were matched to transcription factors whose roles in the binding regulation have not been previously characterized.

The generative nature of the GERV model scores each variant as the predicted change to a proximal ChIP-seq signal. The analysis on six transcription factors NF-*κ*B, CTCF, FOS, JUND, MAX and MYC demonstrated that GERV outperforms existing methods in discriminating variants known to alter TF binding from negative control sets. In a few cases (Figure 3F, Supplementary Figure S4F), the discriminative nature of the competing methods equipped them with higher precision for recalling a small fraction of positives. However their inability to model auxiliary sequences led to the dramatic precision decrease afterwards, while GERV achieved constantly high precision for larger recall.

Applied to an associated variant set (AVS) of breast cancer, GERV correctly predicted the effect of previous validated causal SNP rs4784227, and highly prioritized variants reported to affect FOXA1 binding in breast cancer cell line. With the superior performance exemplified in this task, we expect GERV to play an important role in functionally annotating and prioritizing putative causal variants for downstream experimental analysis

## Acknowledgement

We thank Yuchun Guo for technical support in co-factor analysis using GEM. We also thank Matthew Edwards for many helpful comments and discussions.

